# The strange role of brain lesion size in cognitive neuropsychology

**DOI:** 10.1101/2021.03.27.437336

**Authors:** Christoph Sperber

## Abstract

The size of brain lesions is a variable that is frequently considered in cognitive neuropsychology. In particular, lesion-deficit inference studies often control for lesion size, and the association of lesion size with post-stroke cognitive deficits and its predictive value are studied. In the present article, the role of lesion size in cognitive deficits and its computational or design-wise consideration is discussed and questioned. First, I argue that the commonly discussed role or effect of lesion size in cognitive deficits eludes us. A generally valid understanding of the causal relation of lesion size, lesion location, and cognitive deficits is unachievable. Second, founded on the theory of covariate control, I argue that lesion size control is no valid covariate control. Instead, it is identified as a procedure with only situational benefits, which is supported by empirical data. This theoretical background is used to suggest possible research practices in lesion-deficit inference, post-stroke outcome prediction, and behavioural studies. Last, control for lesion size is put into a bigger methodological and also historical context – it is identified to relate to a long-known association problem in neuropsychology, which was previously discussed from the perspectives of a mislocalisation in lesion-deficit mapping and the symptom complex approach.

Highlights
- Lesion size is a factor that is often considered or controlled in neuropsychology
- No general causal relation between lesion size, lesion location and deficit exists
- Lesion size in brain mapping, outcome prediction and behavioural study is discussed
- Lesion size control is no valid covariate control
- Practical suggestions and guidelines how to consider lesion size are provided

## 1 Introduction

The study of human cognition in brain pathology bears outstanding epistemological value. What it lacks in the ability to uncover temporal aspects of human cognition it compensates with unique advantages, including the ability to infer the existence of separate cognitive processes (Shallice, 1988), the ability to identify anatomical structures that are necessary for a function (Rorden and Karnath, 2004), and insights into how the structural connectome subserves cognition (Catani and ffytche, 2005). However, the epistemological value of human brain lesion-deficit studies also suffers a major limitation. They are purely observational studies in a highly complex, high-dimensional system, and thus any inference of causal relations is a challenge.

A variable that has been thoroughly considered to master this challenge and allow a better understanding of post-stroke deficits is lesion size. In some studies, researchers took lesion size into account either computationally or within their study design to investigate the anatomy of cognitive deficits (e.g., Basso et al., 1987; Karnath et al., 2004; Kalénine et al., 2010; Schwartz et al., 2011; Schwartz et al., 2012; Zhang et al., 2014; Mirman et al., 2015; Winder et al., 2016; Shahid et al., 2017; Sperber and Karnath, 2017; DeMarco and Turkeltaub, 2018; Gajardo-Vidal et al., 2018; Dickens et al., 2019; Garcea et al., 2020). However, this consideration of lesion size has also been criticised (Nachev, 2015; Xu et al., 2018), and the use of lesion size as a covariate should be well-justified given the potential misapplication of covariate control in lesion mapping studies (Sperber et al., 2020). In other studies, researchers directly investigated the role of lesion size in cognitive or behavioural pathology (e.g., Kertesz and Ferro, 1984; Thye and Mirman, 2018). This also includes the use of lesion size as a clinical prognostic factor, most often in combination with other variables (e.g., Goldenberg and Spatt, 1994; Zhu et al., 2010; Long et al., 2011; Kaczmarczyk et al., 2012; Hope et al., 2013; Kuceyeski et al, 2016; Laredo et al., 2018; Loughnan et al., 2019; Ri et al., 2020).

The present article aims to deepen our understanding of the variable lesion size in cognitive neuropsychology and to scrutinise the methodology to consider lesion size in different research areas. First, the relation between lesion size, lesion location and cognitive deficits is discussed (section 3:’*The role of lesion size in cognitive deficits’*). Next, this understanding is used to critically question the strategies to consider lesion size in anatomo-behavioural studies. Likewise, the consideration of lesion size in purely behavioural studies and stroke outcome prediction are discussed (section 4’*The consideration of lesion size in cognitive neuropsychology’*). This technical and detailed section is followed by a concise, simple summary and practical suggestions on the consideration of lesion size in future studies (section 5’ *Conclusions– should we consider lesion size in anatomical or cognitive studies?*’).

## 2 General methods and data availability

Empirical examples and illustrations are based on a publicly available dataset from the Moss Rehabilitation Research Institute at the University of Pennsylvania available in the LESYMAP software package (https://github.com/dorianps/LESYMAP). This dataset includes 131 lesion maps of left hemisphere stroke normalised to MNI space. For more information, see Pustina et al. (2018). Analyses in the present paper were performed with MATLAB 2020a, SPM12 (https://www.fil.ion.ucl.ac.uk/spm/), and NiiStat (https://www.nitrc.org/projects/niistat/).

All data underlying shown examples, scripts, and topographies are publicly available at http://dx.doi.org/10.17632/ff49cy9bbs.1.

## 3 The role of lesion size in cognitive deficits

### 3.1 An intuitive understanding of the role of lesion size

To evaluate the consideration of lesion size in cognitive neuropsychology a detailed understanding of the relation between lesion size, lesion location, and cognitive deficits is required. The previous literature provides a somehow vague picture of this issue. For example, it is stated that “lesion size impacts on executive function” (Long et al., 2011), the “prognostic consequences of lesion size” are studied (Laredo et al., 2018), and mapping studies are supposed to consider the “effects of lesion size” (Thye and Mirman, 2018). This wording hints at a causal role of lesion size in cognitive pathology. But only rarely, this putative causal relation between lesion size and a cognitive deficit is specified in detail (e.g. Karnath et al., 2004; DeMarco and Turkeltaub, 2018; Thye and Mirman, 2018): larger lesions generally induce more and more severe deficits, and they also affect more brain regions. Therefore, a larger lesion is more likely to affect brain structures critical for a deficit and to induce a deficit. To better understand and evaluate these relations by intuition, we can resort to a simple analogy from hunting (Figure 1). Shooting at a duck is more likely to hit and kill the duck if the shot contains more shotgun pellets and covers a larger area.

**Figure 1.**
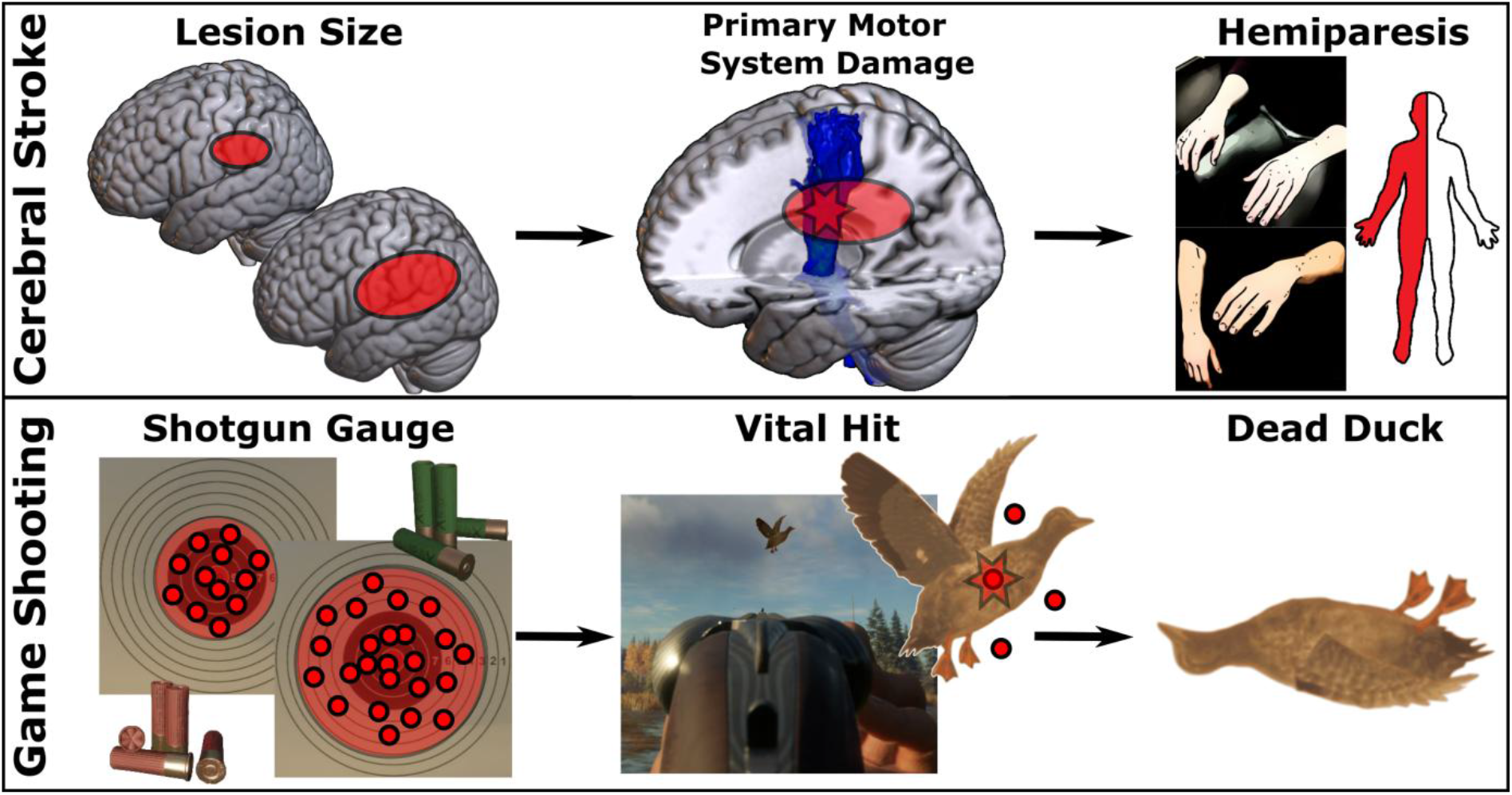
An analogy between lesion size in neurological deficits and hunting.

If we accept this analogy, we can identify a non-deterministic causal relation between shot spread and the duck’s death, but we would not consider shot spread if we merely wanted to understand the relationship between a vital hit and the duck’s death, because shotgun pellet spread plays no role in between these two variables. Intuitively, we would probably tell the whole story of causality – why did the poor duck die? – without referring to the effect of shotgun pellet spread. We aimed, pulled the trigger, fired a shot at the duck, and independent of shot properties, a vital hit is necessary and sufficient to kill the duck. We would also not rely on shot properties if we wanted to predict if a duck survives a bullet wound. In conclusion, the analogy highlights how difficult it is to pinpoint the role of lesion size for cognitive deficits. However, we are not even there yet - this perspective on lesion size falls far too short of reality.

### 3.2 A deeper understanding of the role of lesion size

Cerebral stroke follows the vasculature and has a typical anatomy. We can obtain a rough idea of this typical anatomy with an example data set of 131 normalised lesions (Figure 2A). The assumed non-deterministic causal relation between lesion size and a deficit proves to be even more difficult to grasp if we look at how this anatomy relates to lesion size. First, let us consider the average lesion size by which a voxel is affected. For each voxel lesioned in at least 5 patients, I identified all patients with a lesion including this voxel and computed the average size of their lesions (Figure 2B). This topography highlights that average lesion size massively varies across the brain with a range of 48.3cm³ to 321.5 cm³. For example, lesions to frontal and anterior areas are often large, and lesions to posterior and subcortical areas tend to be much smaller. Second, let us consider the correlation between lesion size and lesion location. For each voxel lesioned in at least 5 patients, I computed across all 131 patients the point-biserial correlation between the binary voxel status (intact/lesioned) and the patient’s lesion size (Figure 2C). In most areas of the brain, the correlation was high with a maximum ρ = 0.75. Hence, this analysis verified the general tendency of voxel-wise damage status to correlate with lesion size. However, this was not the case for all brain regions. Quite the contrary, lesion size and voxel-wise damage status correlated negatively in the caudate and posterior brain areas, and only minimally positively in superior parietal and other subcortical areas. This finding also corresponds to actual deficits that negatively correlate with lesion size, and which are suspected to arise after damage to these very areas (Martinelli et al., 2020). Given these findings, we can re-evaluate the previous perspective on lesion size. A larger lesion is often more likely to affect a brain region that is critical for a cognitive function, but sometimes this relation does not exist or is even inverted. The non-deterministic causal relationship between lesion size and cognitive deficits does not always exist.

**Figure 2.**
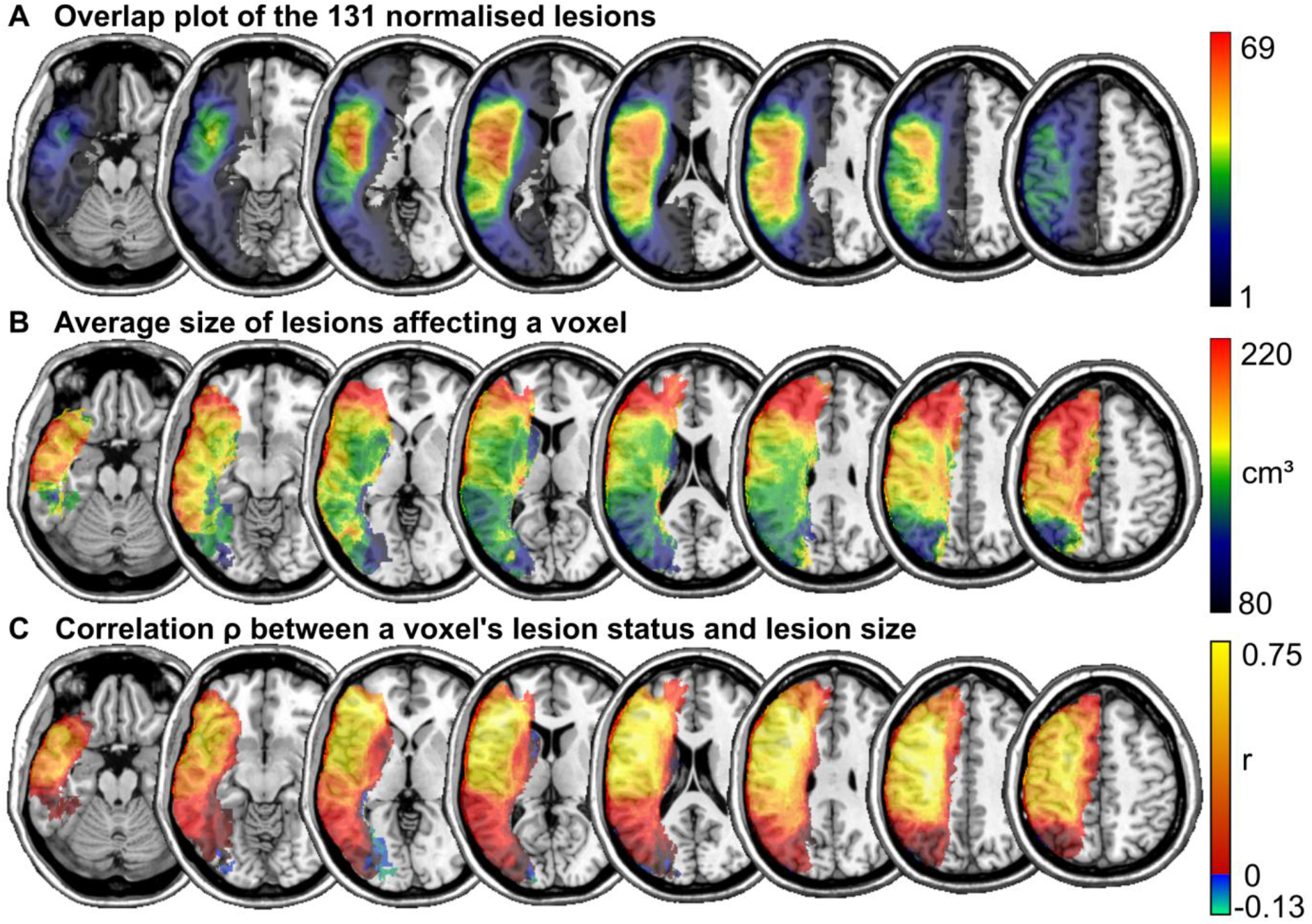
Mapping the complexity of lesion size. Topographies based on the 131 lesions mapping A) the overlap of all lesions, B) the average size of all lesions affecting each voxel, and C) the point-biserial correlation between a voxel’s binary damage status and a patient’s lesion size.

Yet, the situation is still severely oversimplified. So far, we looked at topographies that mapped lesion-size related information on a voxel-wise level. However, the brain is not organised in single voxels. Instead, most cognitive functions are assumed to be subserved by complex networks of connected brain regions. Imagine we want to map the anatomy of a deficit. As it is the nature of a mapping study, we do not know in advance what the deficit’s neural correlates are. It might be a fronto-occipital brain network, including i) occipital regions where lesions are rare and lesion size is negatively correlated with damage, ii) a white matter tract ranging through areas where the correlation between lesion size and damage fluctuates between zero and low positive values, and where relatively small damage can induce a complete disconnection of relevant brain areas with a high impact on behaviour, and iii) a frontal area, where lesion size and damage status is highly correlated. Likewise, the neural correlates might look entirely different. Now, what knowledge can we extract from the fact that lesion size and a deficit moderately correlate by r = 0.3, and what can we conclude about the role of lesion size in this deficit? Not much, if anything at all. The functional anatomy of the brain and the anatomy of stroke appear too complex to allow any definite, generally valid statement about the relationship between lesion size and deficits. Most importantly, it seems impossible to exactly define a causal relationship between both variables, and the commonly used terminology that vaguely implies causality appears to be ill-placed. Lesion size does not appear to have a specific role in, nor an impact or an effect on cognitive deficits. Instead, lesion size appears to be one among a myriad of variables that can be derived from lesion imaging, and it correlates with many variables, including many cognitive deficits.

## 4 The consideration of lesion size in cognitive neuropsychology

### 4.1 Lesion size control in lesion-deficit inference

The role of lesion size and its consideration has been discussed intensely in the context of lesion-behaviour mapping. Already shortly after voxel-wise statistical parametric mapping became popular for lesion-deficit inference (Bates et al., 2003), lesion size was assumed to be linked to a possible bias of topographical results. This bias might be accounted for by covariate control (Karnath et al., 2004). Different kinds of covariate control for lesion size have been used regularly (see DeMarco and Turkeltaub [2018] for an overview on computational approaches) and were also implemented for some multivariate approaches to lesion-deficit inference (Zhang et al., 2014; DeMarco and Turkeltaub, 2018). The putative bias of lesion-deficit topographies related to lesion size was described as a spill-over effect of statistical results, i.e. the appearance of false alarms (Karnath et al., 2004; DeMarco and Turkeltaub, 2018; Thye and Mirman, 2018). This can be explained by two assumptions: i) deficits and lesion size are often positively correlated; ii) in many brain areas, damage status and lesion size are positively correlated (see Figure 2C). *Hence, a deficit will often, at least to a small degree, also correlate with the damage status in many areas of the brain, even if these areas are not the deficit’s neural correlates*.

Interestingly, this description of a spill-over effect perfectly resembles an only much later described bias of topographical results in univariate voxel-based lesion-behaviour mapping (Inoue 2014; Mah et al., 2014). This bias results from the systematic anatomy of lesions (Mah et al., 2014; Nachev, 2015) and can be explained by looking at the collaterality of lesion status between brain regions (Sperber, 2020). Stroke has a typical anatomy and many areas in the brain are often damaged together. If we imagine that a cognitive deficit results from damage to only a small focal area, many areas in the brain are systematically damaged together with this area. *Hence, a deficit will often, at least to a small degree, also correlate with the damage status in many areas of the brain, even if these areas are not the deficit’s neural correlates*.

Given the high concordance of both methodological issues, I argue that researchers described the very same issue from different perspectives. Further, I believe that neither perspective is linked to a methodology that provides a perfect solution. Let us first look at the issue from the perspective of the anatomical bias in univariate lesion-behaviour mapping. The implementation of multivariate lesion-behaviour mapping (MLBM) has been suggested and embraced by large parts of the scientific community to overcome this bias (Mah et al., 2014; Adolphs, 2016; DeMarco and Turkeltaub, 2018). However, empirical results did not confirm the ability of MLBM to overcome it (Sperber et al., 2019; Ivanova et al., 2021), and in a previous publication, I argued that the underlying issue is a problem of causal inference that cannot be solved by replacing associational univariate models with associational multivariate models (Sperber, 2020). Let us now look at the issue from the perspective of a lesion size-related bias. Control for lesion size has been empirically tested by simulations and was found to improve the precision of univariate (Sperber and Karnath, 2017) and multivariate lesion mapping methods (Zhang et al., 2014; DeMarco and Turkeltaub, 2018). Further, control of lesion size was even suggested to partially counteract the anatomical bias in univariate lesion-behaviour mapping (Sperber & Karnath, 2017). Does this mean that lesion size control indeed solves the issues in topographical lesion-deficit inference which putatively require causal inference? Unfortunately, my previous work on this topic (Sperber & Karnath, 2017) meanwhile appears to be theoretically short-sighted to me, and currently, I am convinced that control for lesion size does not provide a good solution. The explanation for my conclusion is rather theoretical: covariate control for lesion size simply does not exist.

#### 4.1.1. Control for lesion size is no valid covariate control

A critical evaluation of covariate control for lesion size requires us to understand two points. First, control for a covariate can be invalid and bias our results. This becomes intuitively evident when looking at a simple experiment (see Figure 3A). Imagine we want to test if the heat of a candle’s flame induces pain. For several trials, we randomly alternate the candle’s state between heat-producing (lit) and non-heat-producing (unlit) and measure a participant’s pain sensation when touching the candle’s end. There will likely be a strong relation between heat and pain. However, heat will also correlate with light. Shall we apply covariate control to remove the variance explained by light from the variable heat? Well, both variables perfectly correlate, and thus controlling for light will remove all variance from the variable heat, and the association between heat and pain will not be present anymore. This simple example shows that covariate control generally can be invalid and misguide our judgements about causal relations in the world. We also see that pure knowledge about the correlation between a dependent variable and the desired covariate variable is not sufficient to identify valid covariate control. Accordingly, it was shown that inappropriate covariate control can grossly affect topographical results in lesion-behaviour mapping (Sperber et al., 2020), and it was previously acknowledged that control for lesion size can also negatively affect results under certain circumstances (Karnath et al., 2004).

**Figure 3.**
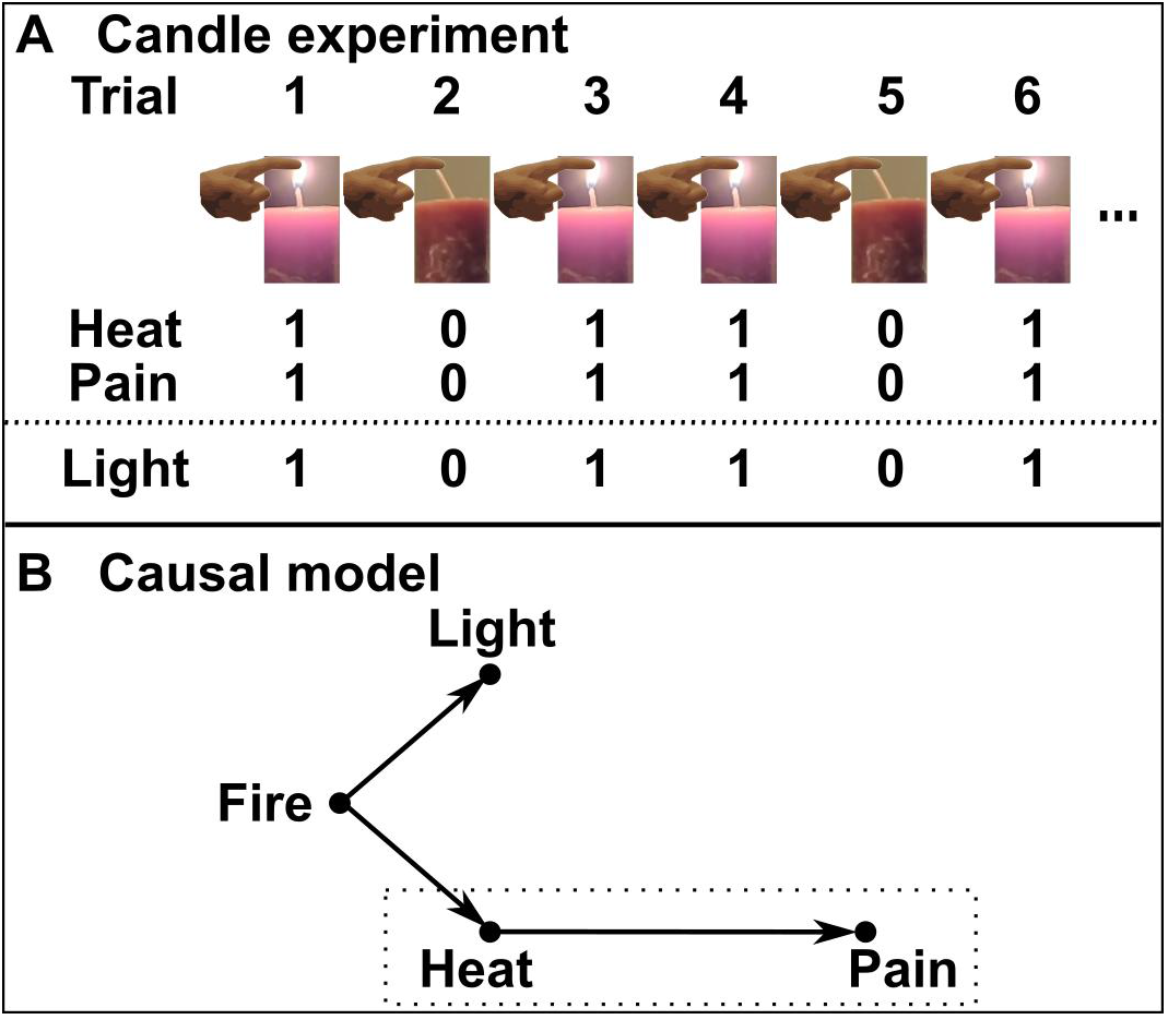
How to identify valid covariate control. A) A simple experiment to test whether a candle’s heat induces pain. We find a perfect relation between heat and pain, but also between pain and light. Should we, therefore, control for light? If we do so, all variance will be removed, because light and heat share the same variance, and the relation between heat and pain will not be present anymore. Obviously, covariate control is invalid here. B) The strategy to identify proper covariates requires us to hypothesize the direction of causal relations between the variables. When we want to infer the causal relation between heat and pain, we should only control for light if there is also a presumed direct causal relation between light and pain, which fulfils the so-called back-door criterion (Pearl & Mackenzie, 2018). As this is not the case, we can formally show that covariate control is invalid here.

The second point that we need to understand is the principle behind the strategy to identify appropriate versus inappropriate covariate control. This requires an evaluation of the presumed causal relations between involved variables (Pearl & Mackenzie, 2018). For example, we can visualise the presumed causal relations for the candle example (Figure 3B) to conclude that covariate control is inappropriate here (for the detailed strategy, see Pearl, 1996; Pearl & Mackenzie, 2018).

Now, what about lesion size control? Whatever exact conditions for causal relations justify covariate control, we will fail to verify them for the factor lesion size. As discussed in length in section 3 (‘*The role of lesion size in cognitive deficits’*), we are unable to identify any generally existing causal relations between lesion size on the one hand and lesion location and cognitive deficits on the other hand. Thus, we are unable to verify lesion size as a covariate that we should control for in the study of lesion-deficit relations. What is commonly termed lesion size control, strictly speaking, is no covariate control.

Still, the consideration of lesion size in lesion-deficit inference was empirically tested and found to improve the quality of lesion-behaviour mapping (Zhang et al., 2014; Sperber and Karnath, 2017; DeMarco and Turkeltaub, 2018). How can this be? What does control for lesion size do if it is no valid control, and why was it found to improve results?

#### 4.1.2. The ambivalence of lesion size control

As described in section 4.1., the problem that researchers aimed to solve by controlling lesion size was to counteract a spill-over effect of significant voxels beyond the actual neural correlates of a function. In the current paper, I argue that lesion size control is not an option to validly control for this spill-over effect. Instead, I argue that, in most cases, lesion size control removed variance, and, by doing so, unspecifically made topographical statistical results more conservative. Therefore, lesion size control has the potential to remove spill-over areas of significant voxels, while it can also do so in areas that belong to a function’s neural correlates. This effect can be intuitively understood by going back to the fictional experiment on the relationship between a flame’s heat and the pain when touching the flame (Figure 3). When mapping pain across several variables (like a deficit across brain image voxels), we find a spill-over effect of results, because pain is not only associated with heat (which caused the pain), but also with light. A control analogue to lesion size control would ideally remove variance until the light-pain association disappears. However, this would, again, not constitute a valid covariate control and the potential benefit of removing spill-over effects also comes as a double-edged sword, as the causal heat-pain association might also disappear. Further, note that the light-pain association does not even constitute a false alarm in the statistical sense, as the non-causal association truly exists.

In lesion-behaviour mapping, such control could be especially problematic if the statistical signal is low, for example, due to a small sample size or a noisy behavioural measure. In this case, the consideration of lesion size might lead to overly conservative results by potentially removing results also from a deficit’s neural correlates, i.e. from areas where the damage caused the symptom. Further, the effect is unspecific. Its intensity might vary depending on the lesion size-deficit association, and there is no way to tell if the consideration of lesion size adequately removed spill-over areas, or if it was too less or too much.

#### 4.1.3. An empirical illustration of the ambivalence of lesion size consideration

For an illustration of the ambivalence of lesion size control, I conducted a simulation experiment. Simulations allow us to empirically test the validity of lesion-behaviour mapping methods. They do not offer high ecological validity, but they perfectly fit an empirical illustration of the ambivalent character of lesion size consideration. The basic principle is that we utilise a sample of real lesions of different sizes following the typical anatomy of stroke. However, we do not investigate true deficits, but simulated deficits. We arbitrarily choose a ground truth area and assign the simulated deficit to all patients with damage to this area. Finally, we map this deficit by lesion-behaviour mapping and evaluate how well the method identifies the ground truth area (for more information, see Sperber & Karnath, 2017). Hence, the trick is that we map deficits for which we have perfect knowledge of their neural correlates, and thus we can evaluate the validity of results.

Thirty different deficits were simulated in the sample of 131 lesions based on regions taken from the BN 246 atlas (Fan et al., 2016). As the lesion-behaviour mapping analysis was computed only for voxels affected in at least 5 patients, voxels with less than 5 lesions were also masked in the atlas. This removed voxels that in any case cannot be correctly identified by lesion-behaviour mapping. Among the remaining atlas regions, the 30 largest were chosen as ground truth regions. Deficits were simulated based on one of these regions each. They were simulated between 0 (no deficit) and 1 (maximal deficit) for each lesion based on the proportion of damaged voxels in the ground truth region. As this procedure alone results in unrealistically high effects (Pustina et al., 2018), I added random noise according to the formula *deficit = (1-x)*plv + x*(rand[0;1])*, whereas *plv* is the proportion of lesioned voxels in the ground truth region, *x* the proportion of noise, and *rand[0;1]* a uniformly random value between 0 and 1. Simulations were performed once for low noise (*x = 10%*), and once for high noise (*x = 50%*). The resulting deficits were mapped by mass-univariate lesion behaviour mapping using general linear models in NiiStat. Sample sizes were varied between the full sample (*N = 131*) and a smaller sample (*N = 50*). Correction for multiple comparisons was applied by maximum statistic permutation correction at p < 0.05. Lesion size was controlled for by the default approach provided by NiiStat, which regresses lesion size out of the target variable. From the resulting topographies, I computed the dependent variables i) ‘false positives’, the amount of significant 1×1×1 mm³ voxels outside of the ground truth area, and ii) ‘sensitivity’, the proportion of voxels inside the ground truth area that are correctly found significant. The statistical analysis was done by a 2×2×2 repeated measures ANOVA for the factors sample size, noise level, and lesion size control; the interpretation and post-hoc tests were focused on effects involving lesion size.

The results are shown in Figure 4. A significant three-way interaction was found for false alarms (F(1,29) = 23.935; p < 0.001). Bonferroni-corrected post-hoc tests found the number of false positives to significantly differ between present vs. absent lesion size control across all combinations of the factors sample size and noise. For sensitivity, the most complex significant term including the factor lesion size was an interaction between lesion size and noise (F(1,29) = 37.112; p < 0.001). Post-hoc tests found sensitivity to significantly differ between present versus absent lesion size control for both noise levels.

**Figure 4.**
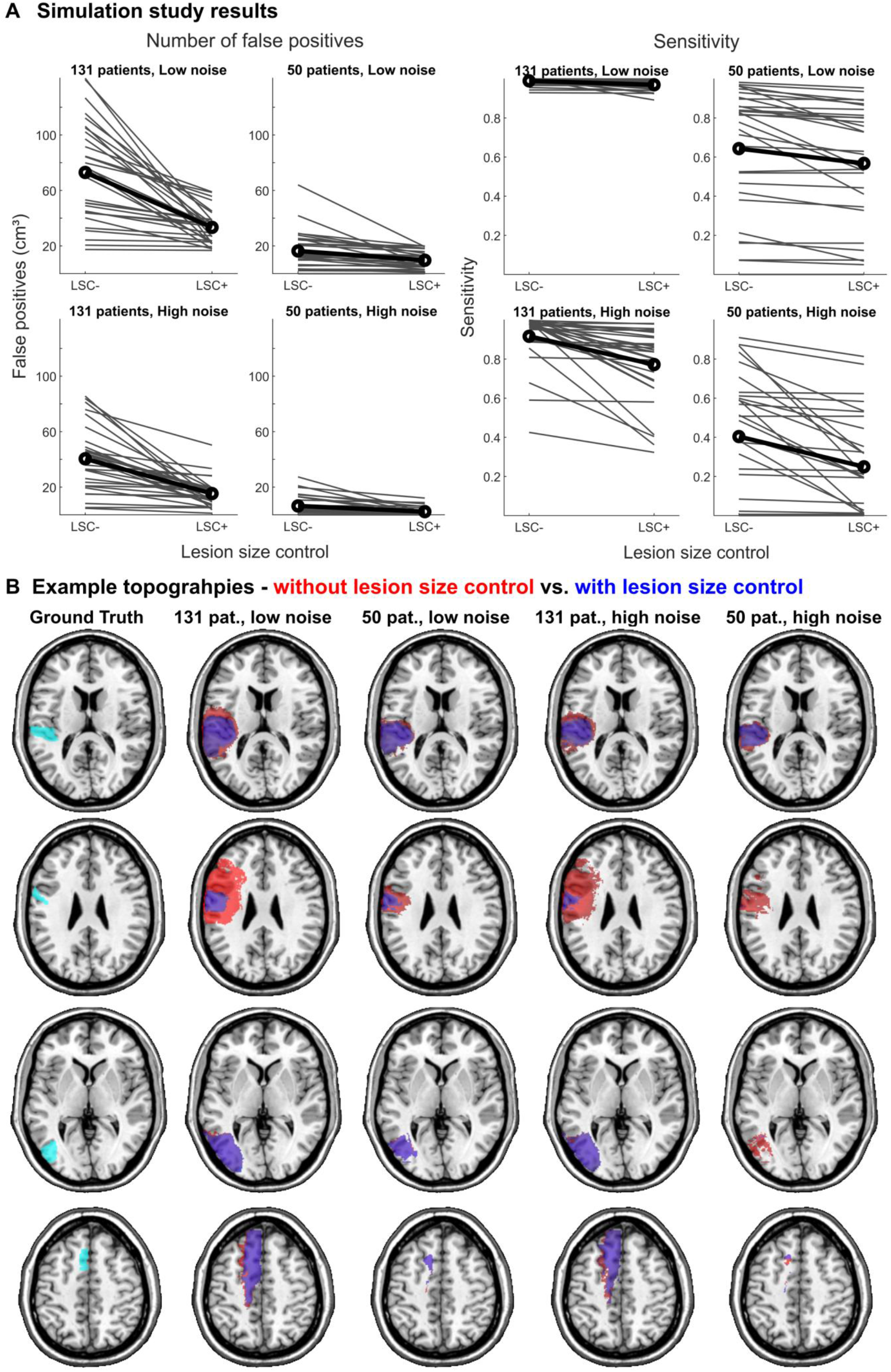
Empirical illustration of the ambivalence of lesion size control. A) Results of the simulation experiment contrasting results without lesion size control (LSC−) with results with lesion size control (LSC+). Bold black lines represent mean values; each of the 30 grey lines represents one single simulation. B) Example topographies for all conditions contrasting results without lesion size control (red) with results with lesion size control (blue).

Hence, the hypothesized ambivalent effects of lesion size control were found in the data. On the one hand, lesion size control reduced the number of false positives and thereby counteracted the spill-over of significant voxels. On the other hand, it reduced sensitivity. Most noticeably, these effects differed across conditions, as indicated by the significant interactions. This is best illustrated by the numerical differences between means for the two most diametral conditions. For the large sample of 131 patients and low noise – the condition that best resembles previous simulation studies (Zhang et al., 2014; Sperber and Karnath, 2017), lesion size control performed favourably with a decrease of false alarms from 73.0cm^3^ to 33.4cm^3^, and a minor decrease in sensitivity from 0.99 to 0.97. For the small sample of 50 patients and high noise – a condition that likely resembles lesion mapping studies with smaller samples – the downside of lesion size control was more apparent, with a decrease of false alarms from 6.4cm^3^ to 2.5cm^3^, and a decrease in sensitivity from 0.40 to 0.25.

This empirical example illustrates several points. First, in line with its postulated ambivalent character, lesion size control did not generally improve results. Hence, the results match the present article’s assumption that lesion size control is no valid covariate control. Second, on a descriptive level, the impact of lesion size control differed markedly across simulations (see grey lines in Figure 4A). This is in accordance with the high variation of the lesion size-deficit association across the brain (Figure 2C). Third, and of the highest practical relevance, the ratio of benefits and drawbacks of lesion size control varied across conditions with different statistical power, i.e. conditions with different sample and effect sizes.

To conclude, the current findings oppose the previous assumptions that a strategy for lesion size control can generally be too anti-conservative (DeMarco and Turkeltaub, 2018). Instead, it seems that the appropriateness of a lesion size control algorithm is situational. Further, even if lesion size control is no valid covariate control, it can be beneficial when a spill-over effect of significant voxels is present. From a purely practical perspective, it would make sense to utilise lesion size control whenever the benefits outweigh the drawbacks. Hence, the need for a strategy to decide if lesion size should be controlled becomes apparent. This is exceptionally difficult in real studies. We can hardly tell how strong the causal lesion-deficit relation is or how reliably the deficit it measured, nor does the mere sample size tell us much about statistical power, as voxel-wise post-hoc statistical power also depends on the lesion anatomy (Kimberg et al., 2007). In the face of these challenges, a possible practical solution is suggested in section 5.

### 4.2 Lesion size in behavioural studies

As outlined in section 3, the association between lesion size and a deficit hardly helps us to gain a deeper understanding of the brain and cognition. Also, while lesion size control has been intensely discussed for topographical lesion-deficit inference, this was much less done so for purely behavioural studies. Does that mean that lesion size is irrelevant here? More specifically we could ask: Is there a behavioural research design where the methodological problems could arise that are described for lesion-behaviour mapping, and why lesion size control is applied?

In lesion-behaviour mapping, lesion size is commonly controlled to counter a spill-over effect of significant lesion-deficit associations beyond a function’s neural correlates, which I argued to be identical with the lesion-anatomical bias of lesion-behaviour mapping (Mah et al., 2014; see section 4.1.). We can go further and not only think of spill-over associations between a deficit with non-causally associated brain areas, but also other deficits that arise after damage to these brain areas. Hence, there could a spill-over of deficit-deficit associations. This is exactly what has been described long ago as the shortcoming of the symptom complex approach (chapter 2.6 in Shallice, 1988, Poeck, 1983), which is well illustrated with what has been termed the Gerstmann syndrome (see Shallice, 1988).

Originally, a complex of the four symptoms pure agraphia, acalculia, finger agnosia, and left-right disorientation was treated as a functional entity, and many theories arose to explain the cognitive architecture that linked these symptoms. Only later, several studies argued that the Gerstmann syndrome does not exist. Instead, Gerstmann described a collection of four symptoms that all arise after parietal damage, i.e. that often co-occur due to proximity of the anatomical correlates and the typical anatomy of lesions.

For this reason, neuropsychology put such a strong focus on behavioural dissociations (Shallice, 1988). Hence, the perspective of the symptom complex approach provides yet another – arguably, also not entirely epistemologically unambiguous – approach to handle a spill-over of associations in neuropsychology. Now, what does the factor lesion size help here? Larger lesions will most often induce more symptoms and the presence of more symptoms will be associated. As in lesion-behaviour mapping, the consideration of lesion size will not help here to obtain incontestable proof. However, lesion size could indicate the relation between symptoms in a group study. An existing syndrome should be identifiable independent of sample characteristics, and especially also in patients with small lesions. Else, if lesion size interacts with the syndrome structure, i.e. how strong symptoms are associated, it seems more likely that the symptoms are independent.

### 4.3 Lesion size in post-stroke outcome prediction

Clinical data assessed in the acute stage of stroke might be used to predict a patient’s potential for recovery after stroke. The only aim in this line of research is to create computational models that optimise predictions. Ironically, our understanding of the human brain and cognition are only of limited use in the creation of good predictors, as modelling strategies to understand functional brain anatomy and to predict outcome from brain imaging data diverge (Bzdok et al., 2020), and imaging features that aim to represent the cause of a deficit can be outperformed by imaging features that do not do so (Sperber et al., 2021). That being said, we can already come to a conclusion: Any understanding of the role of lesion size in cognitive pathology is non-essential here. If lesion size helps to predict outcome, we should use it as a predictor.

Nonetheless, the question remains if lesion size helps predict outcome at all. Does lesion size benefit outcome prediction even after considering lesion location, disconnection (Salvalaggio et al., 2020; Tozlu et al., 2020), functional connectivity (Siegel et al., 2016), demographic and non-imaging clinical data (Price et al., 2017; Tozlu et al., 2020), and so on? This is a question that the present article cannot answer. However, a possible objection can be derived from the correlation analysis on lesion size and damage status shown in Figure 2C. Let us make the arguably unrealistic assumption that lesion size itself genuinely affects and explains post-stroke outcome. Would we then provide prediction models with any new information by adding lesion size after consideration of lesion location, given that lesion location often already highly correlates with lesion size? In other words, do we gain new predictive information from lesion size, if most of its variance is already explained by lesion location? If not, then the addition of lesion size as a predictor will likely be futile.

## 5 Conclusion – should we consider lesion size in cognitive neuropsychology?

The present article argues that although we commonly consider lesion size in cognitive neuropsychology, we are unable to generally define its causal relations with lesion location and the deficit. The role of lesion size in cognitive deficits remains something that can be described by correlations, but a true understanding eludes us. Given that the identification of valid covariate control requires an understanding of the causal relations between variables, it follows that lesion size control is no valid covariate control.

This is a major issue in *lesion-behaviour mapping*, where lesion size control has been used frequently. The rationale behind this control is to remove a spill-over of significant voxels beyond the neural correlates of a deficit. The consequences of invalid covariate control became apparent in an empirical illustration. In some situations, lesion size control mainly removed false positives in the topographical results, while, in others, it mainly decreased the number of true positives. Effect size and sample size appeared to play a central role in this ambivalence of lesion size control. For large samples with strong lesion-deficit relations, i.e. situations where a large spill-over was likely, the benefits prevailed. For smaller samples with medium lesion-deficit relations, the drawbacks prevailed. Hence, even if lesion size control cannot be identified as a valid covariate control, its situation-dependent application appears to be practically useful. Notably, a general rule if lesion size should be controlled for or what control strategy is adequate, and not too conservative or anti-conservative, cannot be stated. Therefore, I suggest the following strategy in lesion-brain inference: In any case, the first result to report should be an uncorrected topography. Next, an a priori criterion to identify a possible spill-over effect should be applied. This criterion could refer, e.g., to the size of the biggest cluster of significant results or the total volume of significant results. For example, we could deem lesion size control appropriate whenever the largest cluster of significant voxels surpasses the size of the largest functional brain region, as defined by the brain atlas used for interpretation. Likewise, a criterion might be generated by reference to previous mapping studies. If the a priori criterion is fulfilled, controlled results are reported, and the main interpretation is based on controlled results.

The specific case of multivariate lesion-behaviour mapping using support vector regression (SVR-LSM; Zhang et al., 2014) deserves a special mention. For this method, and only when using a non-linear machine learning kernel, the consideration of lesion size plays an additional role, as it allows the computation of approximated feature weights. A discussion of this special case can be found in the online supplementary material.

In *behavioural studies*, the role or consideration of lesion size has been less of a topic. However, the interpretation of associations between symptoms was identified as a challenge that suffers from the very same problems as lesion-behaviour mapping. While the consideration of lesion size will not help to obtain proof that human cognition is structured in a certain way, it might indicate certain data interpretations in neuropsychological group studies.

For *post-stroke outcome prediction*, the conclusion is simple and straightforward: if lesion size helps to predict outcome, it should be considered. Only the benefit of its consideration is arguable.

### 5.1 The association problem in neuropsychology

In the present article, I argued that the rationale behind lesion size control traces back to the same methodological challenge in neuropsychology that gave rise to the problem of the symptom complex approach (Shallice, 1988) and the mislocation of lesion-behaviour mapping results (Mah et al., 2014). All these problems originate from the same ambiguity of associations in lesion studies. Arguably, none of these perspectives on the association problem provides an epistemologically perfect solution to unambiguously interpret associations. The consideration of lesion size is no exception here, and its significance in understanding the human brain and cognition might have been overstated in the recent past.

We currently study the brain in times of large data sets, numerous imaging modalities and physiological measurements, and a constantly increasing computational power paired with an ever-growing body of associational multivariate modelling algorithms. If the intention behind this endeavour is to deepen our understanding of the human brain and cognition, we require a critical theoretical groundwork to build the increasing amount of methods upon. A critical, well-reflected consideration of the factor lesion size, or the well-reflected refusal to do so, is a tiny, but valuable step in this journey.

## References

Adolphs, R. (2016). Human Lesion Studies in the 21st Century. Neuron, 90(6), 1151–1153. https://doi.org/10.1016/j.neuron.2016.05.014

Basso, A., Capitani, F., Sala, S. Della Laiacona, M., & Spinnler, H. (1987). Ideomotor apraxia: a study of initial severity. Acta Neurologica Scandinavica, 76(2), 142–146. https://doi.org/10.1111/j.1600-0404.1987.tb03557.x

Bates, E., Wilson, S. M., Saygin, A. P., Dick, F., Sereno, M. I., Knight, R. T., & Dronkers, N. F. (2003). Voxel-based lesion-symptom mapping. Nature Neuroscience, 6(5), 448–450. https://doi.org/10.1038/nn1050

Bzdok, D., Engemann, D., & Thirion, B. (2020). Inference and Prediction Diverge in Biomedicine. Patterns, 1(8), 100119. https://doi.org/10.1016/j.patter.2020.100119

Catani, M., & Ffytche, D. H. (2005). The rises and falls of disconnection syndromes. Brain, 128(10), 2224–2239. https://doi.org/10.1093/brain/awh622

DeMarco, A. T., & Turkeltaub, P. E. (2018). A multivariate lesion symptom mapping toolbox and examination of lesion-volume biases and correction methods in lesion-symptom mapping. Human Brain Mapping, 21(May), 2461–2467. https://doi.org/10.1002/hbm.24289

Dickens, J. V., Fama, M. E., DeMarco, A. T., Lacey, E. H., Friedman, R. B., & Turkeltaub, P. E. (2019). Localization of Phonological and Semantic Contributions to Reading. The Journal of Neuroscience, 39(27), 5361–5368. https://doi.org/10.1523/JNEUROSCI.2707-18.2019

Fan, L., Li, H., Zhuo, J., Zhang, Y., Wang, J., Chen, L., Yang, Z., Chu, C., Xie, S., Laird, A. R., Fox, P. T., Eickhoff, S. B., Yu, C., & Jiang, T. (2016). The Human Brainnetome Atlas: A New Brain Atlas Based on Connectional Architecture. Cerebral Cortex, 26(8), 3508–3526. https://doi.org/10.1093/cercor/bhw157

Gajardo-Vidal, A., Lorca-Puls, D. L., Crinion, J. T., White, J., Seghier, M. L., Leff, A. P., Hope, T. M. H., Ludersdorfer, P., Green, D. W., Bowman, H., & Price, C. J. (2018). How distributed processing produces false negatives in voxel-based lesion-deficit analyses. Neuropsychologia, 115(February), 124–133. https://doi.org/10.1016/j.neuropsychologia.2018.02.025

Garcea, F. E., Greene, C., Grafton, S. T., & Buxbaum, L. J. (2020). Structural Disconnection of the Tool Use Network after Left Hemisphere Stroke Predicts Limb Apraxia Severity. Cerebral Cortex Communications, 1(1), 1–20. https://doi.org/10.1093/texcom/tgaa035

Goldenberg, G., & Spatt, J. (1994). Influence of Size and Site of Cerebral Lesions on Spontaneous Recovery of Aphasia and on Success of Language Therapy. Brain and Language, 47(4), 684–698. https://doi.org/10.1006/brln.1994.1063

Hope, T. M. H., Seghier, M. L., Leff, A. P., & Price, C. J. (2013). Predicting outcome and recovery after stroke with lesions extracted from MRI images. NeuroImage: Clinical, 2(1), 424–433. https://doi.org/10.1016/j.nicl.2013.03.005

Inoue, K., Madhyastha, T., Rudrauf, D., Mehta, S., & Grabowski, T. (2014). What affects detectability of lesion–deficit relationships in lesion studies? NeuroImage: Clinical, 6, 388–397. https://doi.org/10.1016/j.nicl.2014.10.002

Ivanova, M. V., Herron, T. J., Dronkers, N. F., & Baldo, J. V. (2021). An empirical comparison of univariate versus multivariate methods for the analysis of brain–behavior mapping. Human Brain Mapping, 42(4), 1070–1101. https://doi.org/10.1002/hbm.25278

Kaczmarczyk, K., Wit, A., Krawczyk, M., Zaborski, J., & Gajewski, J. (2012). Associations between gait patterns, brain lesion factors and functional recovery in stroke patients. Gait & Posture, 35(2), 214–217. https://doi.org/10.1016/j.gaitpost.2011.09.009

Kalénine, S., Buxbaum, L. J., & Coslett, H. B. (2010). Critical brain regions for action recognition: Lesion symptom mapping in left hemisphere stroke. Brain, 133(11), 3269– 3280. https://doi.org/10.1093/brain/awq210

Karnath, H.-O., Fruhmann Berger, M., Küker, W., & Rorden, C. (2004). The anatomy of spatial neglect based on voxelwise statistical analysis: a study of 140 patients. Cerebral Cortex (New York, N.Y.: 1991), 14(10), 1164–1172. https://doi.org/10.1093/cercor/bhh076

Kertesz, A., & Ferro, J. M. (1984). Lesion size and location in ideomotor apraxia. Brain : A Journal of Neurology, 107 (Pt 3, 921–933. http://www.ncbi.nlm.nih.gov/pubmed/6206911

Kimberg, D. Y., Coslett, H. B., & Schwartz, M. F. (2007). Power in Voxel-based lesion-symptom mapping. Journal of Cognitive Neuroscience, 19(7), 1067–1080. https://doi.org/10.1162/jocn.2007.19.7.1067

Kuceyeski, A., Navi, B. B., Kamel, H., Raj, A., Relkin, N., Toglia, J., Iadecola, C., & O’Dell, M. (2016). Structural connectome disruption at baseline predicts 6-months post-stroke outcome. Human Brain Mapping, 2601(November 2015), 2587–2601. https://doi.org/10.1002/hbm.23198

Laredo, C., Zhao, Y., Rudilosso, S., Renú, A., Pariente, J. C., Chamorro, Á., & Urra, X. (2018). Prognostic Significance of Infarct Size and Location: The Case of Insular Stroke. Scientific Reports, 8(1), 9498. https://doi.org/10.1038/s41598-018-27883-3

Long, B., Anderson, V., Jacobs, R., Mackay, M., Leventer, R., Barnes, C., & Spencer-Smith, M. (2011). Executive function following child stroke: the impact of lesion size. Developmental Neuropsychology, 36(8), 971–987. https://doi.org/10.1080/87565641.2011.581537

Loughnan, R., Lorca-Puls, D. L., Gajardo-Vidal, A., Espejo-Videla, V., Gillebert, C. R., Mantini, D., Price, C. J., & Hope, T. M. H. (2019). Generalizing post-stroke prognoses from research data to clinical data. NeuroImage: Clinical, 24(September), 102005. https://doi.org/10.1016/j.nicl.2019.102005

Mah, Y.-H., Husain, M., Rees, G., & Nachev, P. (2014). Human brain lesion-deficit inference remapped. Brain : A Journal of Neurology, 137(Pt 9), 2522–2531. https://doi.org/10.1093/brain/awu164

Martinelli, F., Perez, C., Caetta, F., Obadia, M., Savatovsky, J., & Chokron, S. (2020). Neuroanatomic correlates of visual hallucinations in poststroke hemianopic patients. Neurology, 94(18), e1885–e1891. https://doi.org/10.1212/WNL.0000000000009366

Mirman, D., Chen, Q., Zhang, Y., Wang, Z., Faseyitan, O. K., Coslett, H. B., & Schwartz, M. F. (2015). Neural organization of spoken language revealed by lesion-symptom mapping. Nature Communications, 6, 6762. https://doi.org/10.1038/ncomms7762

Nachev, P. (2015). The first step in modern lesion-deficit analysis. Brain : A Journal of Neurology, 138(Pt 6), e354. https://doi.org/10.1093/brain/awu275

Pearl, J., Glymour, M., & Jewell, N. P. (2016). Causal inference in statistics (1st ed.). Chichester: Wiley.

Pearl, J., & Mackenzie, D. (2018). The book of why (1st ed.). New York: Basic Books.

Poeck, K. (1983). What do we mean by “aphasic syndromes?” A neurologist’s view. Brain and Language, 20(1), 79–89. https://doi.org/10.1016/0093-934X(83)90034-2

Price, C. J., Hope, T. M., & Seghier, M. L. (2017). Ten problems and solutions when predicting individual outcome from lesion site after stroke. NeuroImage, 145(Pt B), 200– 208. https://doi.org/10.1016/j.neuroimage.2016.08.006

Pustina, D., Avants, B., Faseyitan, O. K., Medaglia, J. D., & Coslett, H. B. (2018). Improved accuracy of lesion to symptom mapping with multivariate sparse canonical correlations. Neuropsychologia, 115(August), 154–166. https://doi.org/10.1016/j.neuropsychologia.2017.08.027

Ri, S., Kivi, A., Urban, P., Wolf, T., & Wissel, J. (2020). Site and size of lesion predict post-stroke spasticity: A retrospective magnetic resonance imaging study. Journal of Rehabilitation Medicine, 52(5). https://doi.org/10.2340/16501977-2665

Rorden, C., & Karnath, H. O. (2004). Using human brain lesions to infer function: A relic from a past era in the fMRI age? Nature Reviews Neuroscience, 5(10), 812–819. https://doi.org/10.1038/nrn1521

Salvalaggio, A., De Filippo De Grazia, M., Zorzi, M., Thiebaut de Schotten, M., & Corbetta, M. (2020). Post-stroke deficit prediction from lesion and indirect structural and functional disconnection. Brain : A Journal of Neurology, 143(7), 2173–2188. https://doi.org/10.1093/brain/awaa156

Schwartz, M. F., Kimberg, D. Y., Walker, G. M., Brecher, A., Faseyitan, O. K., Dell, G. S., Mirman, D., & Coslett, H. B. (2011). Neuroanatomical dissociation for taxonomic and thematic knowledge in the human brain. Proceedings of the National Academy of Sciences, 108(20), 8520–8524. https://doi.org/10.1073/pnas.1014935108

Schwartz, M. F., Faseyitan, O., Kim, J., & Coslett, H. B. (2012). The dorsal stream contribution to phonological retrieval in object naming. Brain, 135(12), 3799–3814. https://doi.org/10.1093/brain/aws300

Shahid, H., Sebastian, R., Schnur, T. T., Hanayik, T., Wright, A., Tippett, D. C., Fridriksson, J., Rorden, C., & Hillis, A. E. (2017). Important considerations in lesion-symptom mapping: Illustrations from studies of word comprehension. Human Brain Mapping, 38(6), 2990–3000. https://doi.org/10.1002/hbm.23567

Shallice, T. (1988). From Neuropsychology to Mental Structure. Cambridge: Cambridge University Press.

Siegel, J. S., Ramsey, L. E., Snyder, A. Z., Metcalf, N. V, Chacko, R. V, Weinberger, K., Baldassarre, A., Hacker, C. D., Shulman, G. L., & Corbetta, M. (2016). Disruptions of network connectivity predict impairment in multiple behavioral domains after stroke. Proceedings of the National Academy of Sciences of the United States of America, 113(30), E4367–76. https://doi.org/10.1073/pnas.1521083113

Sperber, C. (2020). Rethinking causality and data complexity in brain lesion-behaviour inference and its implications for lesion-behaviour modelling. Cortex, 126, 49–62. https://doi.org/10.1016/j.cortex.2020.01.004

Sperber, C., & Karnath, H.-O. (2017). Impact of correction factors in human brain lesion-behavior inference. Human Brain Mapping, 38(3), 1692–1701. https://doi.org/10.1002/hbm.23490

Sperber, C., & Karnath, H.-O. (2018). On the validity of lesion-behaviour mapping methods. Neuropsychologia, 115(May), 17–24. https://doi.org/10.1016/j.neuropsychologia.2017.07.035

Sperber, C., Nolingberg, C., & Karnath, H. (2020). Post‐stroke cognitive deficits rarely come alone: Handling co‐morbidity in lesion‐behaviour mapping. Human Brain Mapping, 41(6), 1387–1399. https://doi.org/10.1002/hbm.24885

Sperber, C., Rennig, J., & Karnath, H. O. (2021). Imaging biomarkers for motor outcome after stroke – should we include information from beyond the primary motor system? bioRxiv 2020.07.20.212175; doi: https://doi.org/10.1101/2020.07.20.212175

Sperber, C., Wiesen, D., & Karnath, H. O. (2019). An empirical evaluation of multivariate lesion behaviour mapping using support vector regression. Human Brain Mapping, 40(5), 1381–1390. https://doi.org/10.1002/hbm.24476

Thye, M., & Mirman, D. (2018). Relative contributions of lesion location and lesion size to predictions of varied language deficits in post-stroke aphasia. NeuroImage: Clinical, 20(June), 1129–1138. https://doi.org/10.1016/j.nicl.2018.10.017

Tozlu, C., Edwards, D., Boes, A., Labar, D., Tsagaris, K. Z., Silverstein, J., Pepper Lane, H., Sabuncu, M. R., Liu, C., & Kuceyeski, A. (2020). Machine Learning Methods Predict Individual Upper-Limb Motor Impairment Following Therapy in Chronic Stroke. Neurorehabilitation and Neural Repair, 154596832090979. https://doi.org/10.1177/1545968320909796

Winder, K., Seifert, F., Köhrmann, M., Crodel, C., Kloska, S., Dörfler, A., Hösl, K. M., Schwab, S., & Hilz, M. J. (2017). Lesion mapping of stroke-related erectile dysfunction. Brain, 140(6), 1706–1717. https://doi.org/10.1093/brain/awx080

Xu, T., Jha, A., & Nachev, P. (2018). The dimensionalities of lesion-deficit mapping. Neuropsychologia, 115(May), 134–141. https://doi.org/10.1016/j.neuropsychologia.2017.09.007

Zhang, Y., Kimberg, D. Y., Coslett, H. B., Schwartz, M. F., & Wang, Z. (2014). Multivariate lesion-symptom mapping using support vector regression. Human Brain Mapping, 35(12), 5861–5876. https://doi.org/10.1002/hbm.22590

Zhu, L. L., Lindenberg, R., Alexander, M. P., & Schlaug, G. (2010). Lesion load of the corticospinal tract predicts motor impairment in chronic stroke. Stroke; a Journal of Cerebral Circulation, 41(5), 910–915. https://doi.org/10.1161/STROKEAHA.109.577023

