## Supplementary text for "The strange role of brain lesion size in cognitive neuropsychology"

Christoph Sperber

*The consideration of lesion size in SVR-LSM*

The consideration of lesion size in support vector regression-based lesion-symptom mapping (SVR-LSM; Zhang et al., 2014), a multivariate approach to lesion-behaviour mapping, deserves to be dealt with separately. The conclusions drawn in section 4.1. should also apply here. However, another specific issue arises for SVR-LSM using a support vector regression (SVR) with radial basis function (RBF) kernel, which is the default kernel in most software packages (Zhang et al., 2014; DeMarco and Turkeltaub, 2018). SVR-LSM requires the computation of a feature weight for each voxel (or other input feature, respectively). Feature weights can only be computed in an SVR with a linear kernel, but not in an SVR with a non-linear RBF kernel. However, Zhang et al. (2014, see appendix section there) found an approximation that allows the computation of feature weights in SVR with RBF kernel. This approximation assumes that some mathematical terms become asymptotically irrelevant due to input features – the voxel-wise lesion data – becoming vanishingly small after a certain kind of lesion size correction. So-called direct total lesion volume control (dtlvc) transforms binary 1/0 lesion maps into binary (1/lesion size in voxel)/0 lesion maps, i.e. lesion maps where the lesion area is represented binarily with a number much smaller than 1. If one accepts this approximation, feature weights can be computed for SVR with an RBF kernel the same way as for SVR with a linear kernel. Still, the question remains if a violation of this approximation invalidates SVR-LSM. Other strategies to consider lesion size in SVR-LSM have been used (DeMarco and Turkeltaub, 2018). While results for actual behavioural deficits appear overly conservative (see Table 2 in DeMarco and Turkeltaub, 2018), results under well-controlled simulation condition perform partially even better than under direct total lesion volume control (see Figure 6 in DeMarco and Turkeltaub, 2018). In conclusion, the optimal way to perform SVR-LSM remains an open question, and there might be some computational pitfalls related to the way lesion size is controlled.

**References**

DeMarco, A. T., & Turkeltaub, P. E. (2018). A multivariate lesion symptom mapping toolbox and examination of lesion-volume biases and correction methods in lesion-symptom mapping. *Human Brain Mapping*, *39*(11), 4169–4182. https://doi.org/10.1002/hbm.24289

Zhang, Y., Kimberg, D. Y., Coslett, H. B., Schwartz, M. F., & Wang, Z. (2014). Multivariate lesion-symptom mapping using support vector regression. *Human Brain Mapping*, *35*(12), 5861–5876. https://doi.org/10.1002/hbm.22590
